# Practice beyond performance stabilization increases the use of online adjustments to unpredictable perturbations in an interceptive task

**DOI:** 10.1101/2024.10.30.621165

**Authors:** Crislaine Rangel Couto, Cláudio Manoel Ferreira Leite, Carlos Eduardo Campos, Leonardo Luiz Portes, Cíntia de Oliveira Matos, Suziane Peixoto Santos, Natália Fontes Alves Ambrósio, Hani Camille Yehia, Herbert Ugrinowitsch

**Affiliations:** Sports Departament, Universidade Federal de Minas Gerais-UFMG, Belo Horizonte, Minas Gerais, Brazil; Departament of Eletronic Enginnering, Universidade Federal de Minas Gerais-UFMG, Belo Horizonte, Minas Gerais, Brazil; Departament of Physical Education and Health Science, Universidade Federal de São João del-Rei-UFSJ, São João del-Rei, Minas Gerais, Brazil; Department of Mathematics and Statistics, University of Western, Perth, Western Australia, Australia; Departament of Sports Science, Universidade Federal do Triângulo Mineiro-UFTM, Uberaba, Minas Gerais, Brazil; Department of Prevention and Health Promotion, Centro Universitário Una, Belo Horizonte, Minas Gerais, Brazil

**Keywords:** Stabilization level, Superstabilization level, Motor control, Perturbations, Pre-programation, Feedback online

## Abstract

In recent decades, research has focused on motor adjustments in interception tasks within predictable environments. However, emerging studies suggest that continued practice beyond performance stabilization enhances the ability to adapt to unpredictable events. The objective of this study was to investigate the effects of practicing until performance stabilization versus extended practice through superstabilization on the ability to adjust to unpredictable perturbations in intercepting a moving target. We hypothesized superstabilization would better facilitate motor adjustments in response to unpredictable perturbations. Forty participants engaged in an interception task until they achieved either performance stabilization or superstabilization. Subsequently, both stabilization and superstabilization groups were tested in an unpredictable environment, where, in certain trials, the target’s velocity unexpectedly changed after the onset of the movement. The findings revealed that the superstabilization group made more adjustments in response to these perturbations than the stabilization group, attributed to their developed capacity to use online feedback as a control mechanism more efficiently. In contrast, the practice until performance stabilization did not foster this adaptive mechanism. These results support the notion that learning is a dynamic process that extends beyond the point of performance stabilization, emphasizing the benefits of continued practice for mastering complex motor tasks in variable contexts.

## Introduction

Hitting a ball during a tennis game requires intercepting a shot from the opposite court. Interception actions of moving targets clearly reveal distinct abilities to solve motor problems due to different skill levels [1]. These differences in skill level become even more apparent when the task is performed in complex and dynamic contexts [2] characterized by environmental changes or perturbations requiring adjustments in an action [3].

Perturbations are a constant in motor performance (e.g., a ball that changes speed after touching the floor) and impose specific sensorimotor demands requiring specific motor adjustments. These perturbations can be predictable or unpredictable to the performer [4, 5]. Predictable perturbations are identified before the movement onset; consequently, the action planning may contain the changes necessary to adapt to the perturbation. Conversely, unpredictable perturbations are identified only after the onset of movement; consequently, the action planning has to be reorganized after its triggering to adapt to the perturbation. In essence, unpredictable perturbations in interception tasks necessitate real-time, online corrections.

Many aspects of motor learning influence the ability to perform online corrections [6] and, consequently, the performance under perturbations [7, 8]. For instance, practice beyond performance stabilization increases the skill level and favors performance in the face of perturbations, even though no apparent difference in performance may occur during practice [3, 9]. Furthermore, a high level of performance stabilization seems to increase accuracy and the sensorimotor system’s high adaptability [3, 10, 11].

Practice until performance stabilization helps build a robust and consistent internal representation of the limbs involved in the task and environmental dynamics [12, 13]. This internal representation of the task allows the performer to use the pre-program mechanism to control the action [10, 14]. However, this pre-programmed mechanism may prove inefficient when faced with an unpredictable perturbation. Incontrast, practice that extends beyond performance stabilization, name superstabilization [3, 9] appears to increase the flexibility of the internal representation [3, 13], thus allowing performers to utilize feedback control mechanisms for making online adjustments.

Flexible internal representations enable efficient motor control by facilitating the extraction and utilization of task-relevant information to generate appropriate adjustments [15, 16]. For example, in coincident timing tasks, expert baseball athletes present a remarkable adaptive capacity in the face of changes in stimulus speed thanks to sophisticated speed detection and response adjustment strategies [16]. Although coincident timing tasks provide an approximation to interceptive actions as these types of tasks present perceptual similarities [17], interceptive actions show some perceptual-motor particularities that require specific consideration, such as an actual approximation of the effector to the moving target and a clear hit/miss condition [18–21] that might influence the mechanisms of control.

The mechanism of control in interceptive motor tasks can be observed from the kinematics analysis, such as time to peak velocity and the number of phases in the acceleration curve of the action [18, 22, 23]. A peak velocity that coincides with or close to the moment of interception, along with a monophasic acceleration curve, typically indicates a predominance of pre-programming mechanism [17, 24]. In these cases, the acceleration curve (velocity derivative) presents a single positive component (i.e., monophasic profile). The same monophasic shape of the acceleration curve has been found when the target travels at a constant speed and predictably changes velocity before the movement onset [19, 20]. Similar results were found when the target accelerates or decelerates [20].

The practice of an interception task in predictable contexts favors the use of the pre-programming mechanism [25]. In such environments, the motor control system can predict the outcomes of actions based on sensory information gleaned from previous practice trials [26, 27]. In contrast, an unstable environment demands a feedback control mechanism, which adjusts the function to environment changes [28, 29]. Furthermore, the successful interception of moving interception demands great prediction and planning to allow sufficient time for adjustments when necessary [30]. For example, Fialho and Tresilian [20] showed that in interceptive tasks, even when the movement time is between 130 and 170 ms regardless of the target velocity, all acceleration curves presented a monophasic profile, demonstrating the need of more time for any adjustment by the motor control system.

In unpredictable contexts, when the target’s speed changes after the onset of the onset, maintaining or achieving proper performance levels requires the use of online feedback mechanisms of control [31]. Under these conditions, the peak of velocity occurs earlier, and the velocity curve presents inflections indicating movement adjustments. For example, Tresilian and Plooy [18] found valleys in the acceleration curve (i.e., bi or polyphasic profile according to the number of valleys), signaling corrective sub-movements during an interception task. The efficiency of the online feedback mechanism in rapid movements hinges on swift corrections to the motor command [32] and seems to be linked to performance stabilization [3, 13]. Although many daily activities and sports involve interception, and the pursuit of success in such tasks is widespread [1], to our knowledge, the relation between the level of performance stabilization level and the mechanisms underlying adjustments to unpredictable perturbations in interceptive actions has yet to be thoroughly investigated.

The purpose of the present study is to investigate the effects of two levels of performance stabilization on the adjustments to unpredictable perturbations in the interception of a moving target. We adopted a virtual interceptive task to manipulate two levels of performance stabilization during the learning phase, followed by a testing phase with unpredictable perturbations interspersed with control trials, i.e., the same condition as the learning phase.

We assumed that practice beyond performance stabilization will enhance online adjustments to unpredictable perturbations, as evidenced by the movement’s kinematics observed on an increased number of phase switches in the acceleration curve and a reduced time to reach peak velocity. This will lead to improved performance under unpredictable conditions.

## Material and Methods

### Participants

Forty-two young university students (26.02 ± 2.02 yr.; 22 men), self-declared right-handed, with a normal or corrected-to-normal vision, no history of neurological impairment or orthopedic limitations of the upper limbs, and inexperienced in the task participated in this study. This sample size was determined by using GPower (version 3.1.2; Franz Faul, Universitat Kiel, Germany) [33]. The following parameters were used to determined the sample size: a power of 0.80; expected effect size of 0.3. Our sample size was set *N* = 18, but a larger sample size was used because this could increase the reability of our results. All participants provided written informed consent prior to testing and were informed that they could withdraw their consent at any time. The local ethics committee approved this study (n. 30544714.7.0000.5149), and the procedures followed the ethical standards in the Declaration of Helsinki 1964, amended in 1989.

### Instruments and Task

The setup for the virtual interception task (Fig.1) consisted of an Intel Celeron 2.20 GHz computer, a 17” monitor (Dell 60 Hz, 1366×768’), a 35 cm long wireless drawing tablet (WACON-INTUOS 3 - 9 x 12) with a capture frequency of 200 Hz, and a digital pen (INTUOS 3). A foaming (EVA) plate 2 cm thick with a posterior-anterior cut in its center (27.7 cm long and 1.7 cm wide) was landed on the tablet, forming a groove constraining the digital pen movement to 1 degree of freedom (i.e., forward). Moreover, the task was projected (SONY VLP ES7® with 60Hz) onto a 304 cm wide and 228 cm high white screen placed 370 cm in front of the participant. The Virtual Interception Task (VIT), developed in Labview® (Leonardo Portes; Crislaine Rangel Couto & Herbert Ugrinowitsch, UFMG, Belo Horizonte/MG, Brazil), controlled the interception task, data acquisition, and processing. The VIT synchronized the virtual task (target and effector) and the tablet and controlled the target velocities.

**Fig 1.**
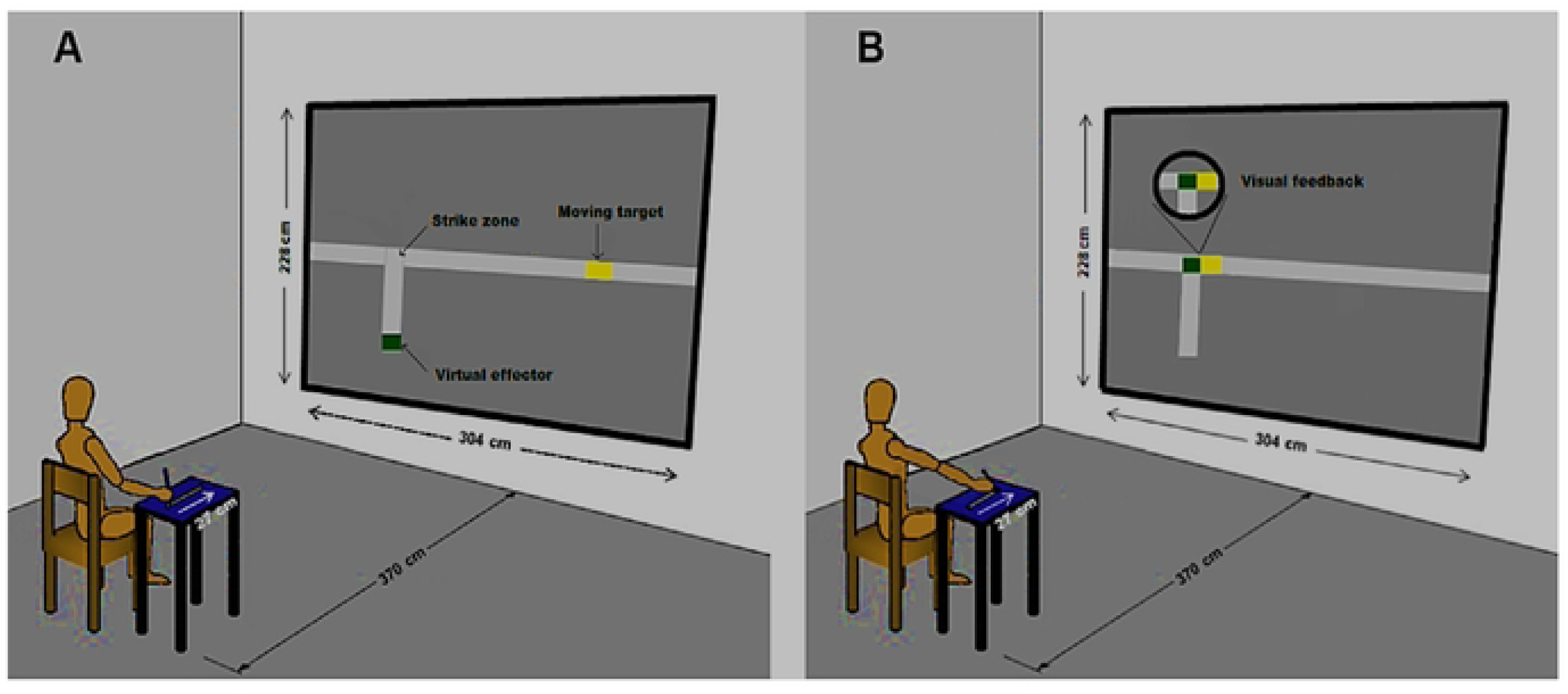
The virtual target (A) moved at a velocity controlled by VIT, over 213 cm to reach the strike zone (B). To control the virtual effector to reach the strike zone at the same moment as the target, the participant moved a pen over the tablet within 200 to 250 ms.

The Interception Task (IT) was to hit a virtual target (4 x 6 cm yellow rectangle) that moved along a horizontal rail 300 cm long, both projected on the screen. To initiate every trial, the participant positioned the pen in the proximal part of the groove (closer to the body), maintaining the elbow joint at 90°. The yellow virtual target appeared on the rightmost side of the screen, and the experimenter controlled the moment the target started traveling through the rail to the left end of the rail, which ranged from 1 to 3 s. When the target was near the interception area, the participant had to move the digital pen along the groove (shoulder flexion and elbow extension). The digital pen controlled the screen’s virtual effector (2 x 4 cm green rectangle), which moved along the vertically projected rail. The participants had to move the virtual green effector and intercept the yellow target within the width of the target and inside the strike zone, which corresponded to intercepting the target within a 5 cm bandwidth relative to its center.

### Procedure and Design

Every participant received demonstrations and plain explanations about the task. Then, he/she sat on a chair with the digital table on the right-hand side. The chair was regulated on the horizontal and vertical axis to maintain the elbow joint at 90° about the shoulder. The target velocity was 145 cm/s, corresponding to a time window of 69 ms, representing the period for which contact with a moving target was possible [34].

The 42 participants were balanced by sex and randomly assigned into two groups previously prepared according to the level of stabilization required for the experiment: a Stabilization Group (SG) and a Superstabilization Group (SSG). This preparation consisted of performing the IT at a constant speed (145 cm/s) until they reached a group-specific performance criterion related to intercepting the target. The Stabilization Group practiced until they intercepted the target three trials in a row. The Superstabilization Group practiced until they reached the same performance criterion (three right trials in a row) six times, which differentiated the Stabilization and Superstabilization groups. As such, the two groups represented two different levels of learning and guaranteed the manipulation of our independent variable. The SG practiced the task until reaching the performance criterion with an error smaller than or equal to 5 cm in 200 trials. The error was calculated by comparing the distance of the effector’s center to the target’s center. The SSG practiced the task until reaching its performance criterion within 320 trials and the same error bandwidth of 5 cm. This criterion follows the procedures of previous studies when two different levels of stabilization showed no difference in performance accuracy when the visual stimulus ran at a constant velocity [3, 9]. Two participants failed to meet the criterion of superstabilization performance over 320 trials and were not included in data analysis.

Besides reaching the established performance criterion, movement time should range from 200 to 250 ms. This duration allowed the participant to pre-program their movements and use online feedback control [18] when necessary. To keep movement duration within the specified range, we provided qualitative verbal information about it after every trial in the following way: (a) Movement times (MTs) below 179ms – “the movement was very fast;” (b) MTs between 180 and 199ms - “the movement was fast;“(c) MTs between 200-250ms - “good movement time;” (d) MTs between 251-270ms - “the movement was slow;” and (e) MTs over 271ms - “the movement was very slow”. Also, the IT projected an image of the green effector and the yellow target at the moment the effector crossed the interception area, which served as a post-trial visual knowledge of results (KR). The participant could request a rest break at any time during the preparation.

After 10 min of reaching the performance criterion started the Exposure phase, consisting of 129 trials. In 111 of them, the target traveled at the same velocity as the Pre-exposure (i.e., 145 cm/s), corresponding to a time window of 55,17 ms. In the other 18 trials, the stimulus started moving at 145 cm/s, but the velocity changed immediately after the onset of the interception movement. These changes randomly increased to 200 cm/s named PI, corresponding to a time window of 40 ms, or decreased to 90 cm/s, named PII, corresponding to a time window of 90 ms. The participants were instructed that the target velocity would change, but they did not know how or in which trials would occur, making perturbations unpredictable (Figure 2).

**Fig 2.**
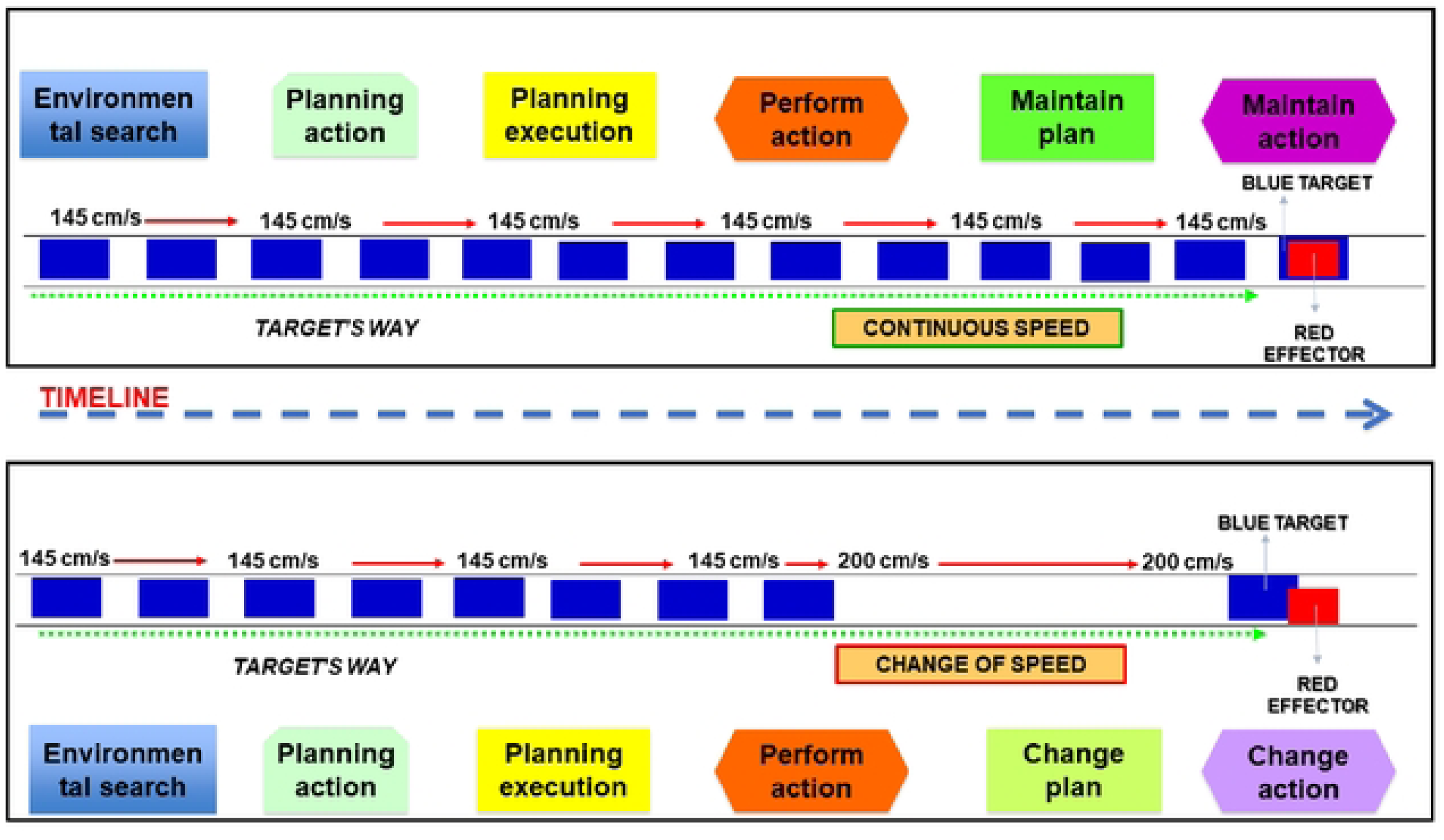
Simulation of the experimental protocol during the exposure phase. The upper part shows a control trial when the target starts moving in the direction of the strike zone at constant velocity. Firstly, the participant searches for environmental information, plans the action, executes the plan, and starts the action. Based on the same target velocity, the participant maintains the action plan and the action. The bottom part shows a perturbation trial when the target changes velocity after the onset of the movement. Firstly, the participant searches for environmental information, plans the action, executes the plan, and starts the action. Based on the change in target velocity, the participant changes the action plan and the action trying to reach the target.

### Measures and data analysis

The constant error (CE), expressed as the magnitude and direction of deviation from the center of the target in cm indicated performance accuracy [35]. The relative time to peak velocity (tPV %), the time to peak velocity expressed as a percentage of the total movement time [36], and the number of corrective sub-movements (N-cor), considered as the number of inflections (valleys) in the acceleration curve [18, 37], indicated the strategies and mechanisms of control. Valleys in the acceleration curve indicated corrective sub-movements when they reached at least 2% of the magnitude of the previous acceleration peak [18].

We analyzed data from the exposition, and we first separated PI and PII and their respective and immediately preceding (Pre) and following (Post) trials because PI and PII should cause opposite error bias. Then, we further divided each perturbation into three particular moments of the exposition (i.e., blocks), indicated by subscript Arabic “algorism”: early exposition (PI_1_ and PII_1_), intermediate exposition (PI_2_ and PII_2_), and late exposition (PI_3_ and PII_3_). Each of these moments was organized in one block of three perturbation trials and their respective pre- and post-trials. This organization analyzed: a) Block 1, with Pre PI_1_, PI and Post PI_1,_ b) Block 2, with Pre PI_2_, PI_2_ and Post PI_2_, and c) Block 3, with Pre PI_3_, PI_3_ and Post PI_3_. PII adopted the same organization. We then obtained the mean value for each variable, CE, tPV %, and N-cor. The CE and tPV% in each moment of the exposition were analyzed by groups (SG and SSG) × blocks of three trials (Pre, P, Post) ANOVA with repeated measures on the blocks factor. To analyze the number of corrections (N-cor), we considered the average number of corrections from the trials with either perturbation. We applied t-Student tests to independent samples to compare the number of corrections performed by SG and SSG during each perturbation (i.e., PI and PII).

The significance level was set at p < 0.05 for all inferential statistics, and Tukey’s post hoc test was used for pairwise comparisons. Additionally, to examine the magnitude of the effects for each analysis, we used partial eta squared (*η_p_*^2^). Effects were considered large when *η_p_*^2^ was above 0.14; medium for *η_p_*^2^ between 0.06 and 0.14; small for *η_p_*^2^ between 0.01 and 0.06; and trivial when *η_p_*^2^ was below 0.01 [38]. Preliminary data analyses indicated that all data fit normality (Shapiro-Wilk’s test), (p > 0.07).

## Results

### Perturbation I (fast velocity)

#### Pre PI_1_ x PI_1_ x Post PI_1_

The analysis of the number of corrections on the first moment of PI (PI_1_) shown in Fig 3 indicates that SSG performed more corrections than SG (*p* = 0.03). The analysis of tPV (%) did not show difference between groups, *F*(1, 38) = 0.27, *p =* 0.60, *η_p_*^2^= 0.007, power (1-β) = 0.08; blocks, *F*(2, 76) = 0.61, *p =* 0.54, *η_p_*^2^= 0.01, power (1-β) = 0.14; nor interaction effects, *F*(2, 76) = 1.23, *p =* 0.29, *η_p_*^2^= 0.03, power (1-β) = 0.26. The analysis of CE showed difference between blocks, *F*(2, 76) = 98,56, *p =* 0.001, *η_p_*^2^= 0.72, power (1-β) = 1.00. The post hoc detected that the CE increased when the perturbation was inserted (PI_1_) compared to Pre PI_1_ (*p <* 0.05). With the withdrawal of perturbation (Post PI_1_), the CE decreased compared to PI_1_ (*p <* 0.05). There was no significant difference between groups, *F*(1, 38) = 0.91, *p =* 0.76, *η_p_*^2^= 0.002, power (1-β) = 0.06 nor main interaction effects, *F*(2, 76) = 2.69, *p =* 0.07, *η_p_*^2^= 0.06, power (1-β) = 0.51.

**Fig 3.**
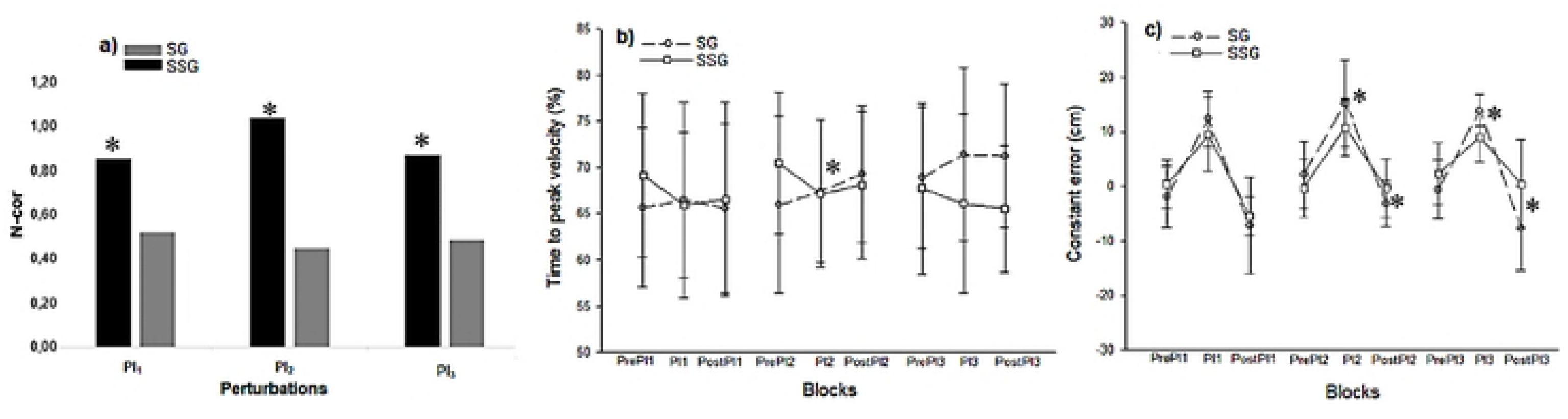
Perturbation I (Fastest Speed). **a)** Mean of the Number of Corrections (N-cor). Black bars represent SSG and gray bars represent SG. **b)** Mean and standard deviation of the time to peak velocity (%) and **c)** mean and standard deviation of the constant error (cm). SG represents stabilization group practice. SSG represents superstabilization group practice. PI represents perturbation with increased velocity. *Interactions differences.

#### Pre PI_2_ x PI_2_ x Post PI_2_

The analysis of the number of corrections on the second moment of PI (PI_2_) showed that SSG performed more corrections than SG (*p* = 0.001). The analysis of tPV (%) showed main interaction effects, *F*(2, 76) = 3.27, *p =* 0.04, *η_p_*^2^= 0.07, power (1-β) = 0.60. The post hoc detected that SSG spent less tPV (%) in PI_2_ than in Pre PI_2_ (*p* < 0.05). There was no significant difference between groups, *F*(1, 38) = 0.21, *p =* 0.64, *η_p_*^2^= 0.005, power (1-β) = 0.07 nor blocks *F*(2, 76) = 0.75, *p =* 0.01, *η_p_*^2^= 0.72, power (1-β) = 0.17. The analysis of CE showed main interaction effects, *F*(2, 76) = 4.94, *p =* 0.009, *η_p_*^2^= 0.11, power (1-β) = 0.79. The post hoc detected that the CE increased when the perturbation was inserted (PI_2_) compared to Pre PI_2_ for both groups (*p <* 0.05). With the withdrawal of perturbation (Post PI_2_), the CE decreased compared to PI_2_ for both groups (*p <* 0.05). There was difference between blocks, *F*(2, 76) = 86,65, *p =* 0.001, *η_p_*^2^= 0.69, power (1-β) = 1.00. The post hoc detected that the CE increased when the perturbation was inserted (PI_2_) compared to Pre PI_2_ (*p <* 0.05). There was no significant difference between groups, *F*(1, 38) = 1.29, *p =* 0.26, *η_p_*^2^= 0.03, power (1-β) = 0.19.

#### Pre PI_3_ x PI_3_ x Post PI_3_

The analysis of the number of corrections on the third moment of PI (PI_3_) showed that SSG performed more corrections than SG (*p* = 0.02). The analysis of tPV (%) did not show any difference between groups, *F*(1, 38) = 2.99, *p =* 0.09, *η_p_*^2^= 0.07, power (1-β) = 0.39; blocks *F*(2, 76) = 0.08, *p =* 0.91, *η_p_*^2^= 0.002, power (1-β) = 0.06; nor main interactions effects, *F*(2, 76) = 2.43, *P =* 0.09, *η_p_*^2^= 0.06, power (1-β) = 0.47. The analysis of CE showed main interaction effects, *F*(2, 76) = 11.99, *p =* 0.001, *η_p_*^2^= 0.23, power (1-β) = 0.99. The post hoc detected that the CE increased when the perturbation was inserted (PI_3_) compared to Pre PI_3_ for both groups (*p <* 0.05). With the withdrawal of perturbation (Post PI_3_), the CE decreased compared to PI_3_ for both groups (*p <* 0.05). There was difference between blocks, *F*(2, 76) = 67.88, *p =* 0.001, *η_p_*^2^= 0.64, power (1-β) = 1.00. The post hoc detected that the CE increased when the perturbation was inserted (PI_3_) compared to Pre PI_3_ (*p <* 0.05). There was no significant difference between groups, *F*(1, 38) = 3.27, *p =* 0.07, *η_p_*^2^= 0.07, power (1-β) = 0.42.

### Perturbation II (slow velocity)

#### Pre PII_1_ x PII_1_ x Post PII_1_

The analysis of the number of corrections on the first moment of PII (PII_1_) shown in Fig 4 indicates that SSG performed more corrections than SG (*p* = 0.01). The analysis of tPV (%) did not show difference between groups *F*(1, 38) = 0.03, *p =* 0.84, *η_p_*^2^= 0.00, power (1-β) = 0.05; blocks, *F*(2, 76) = .70, *p =* 0.49, *η_p_*^2^= 0.01, power (1-β) = 0.16 nor interaction effects, *F*(2, 76) = 0.76, *p =* 0.47, *η_p_*^2^= 0.01, power (1-β) = 0.17. The analysis of CE showed difference between blocks, *F*(2, 76) = 61.05, *p =* 0.001, *η_p_*^2^= 0.61, power (1-β) = 1.00. The post hoc detected that the CE increased when the perturbation was inserted (PII_1_) compared to Pre PII_1_ (*p <* 0.05). With the withdrawal of perturbation (Post PII_1_), the CE decreased compared to PII_1_ (*p <* 0.05). There was no significant difference between groups, *F*(1, 38) = 0.87, *p =* 0.35, *η_p_*^2^= 0.02, power (1-β) = 0.14 nor main interaction effects, *F*(2, 76) = 0.20, *p =* 0.81, *η_p_*^2^= 0.005, power (1-β) = 0.08.

**Fig 4.**
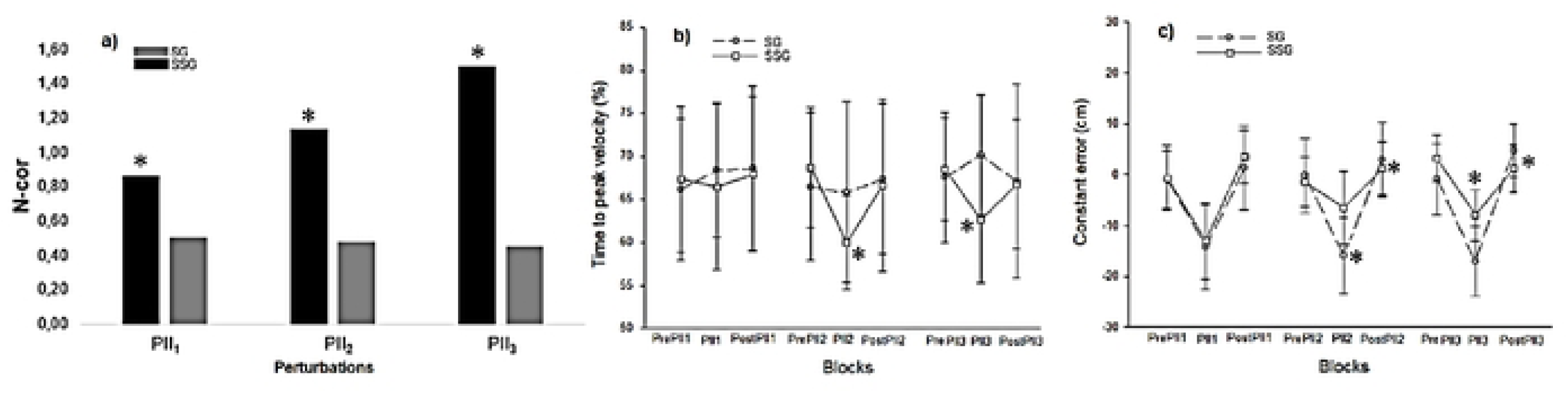
Perturbation II (Slowest Speed). **a)** Mean of the Number of Corrections (N-cor). Black bars represent SSG and gray bars represent SG. **b)** Mean and standard deviation of the time to peak velocity (%) and **c)** mean and standard deviation of the constant error (cm). SG represents stabilization group practice. SSG represents superstabilization group practice. PI represents perturbation with increased velocity. *Interactions differences.

#### Pre PII_2_ x PII_2_ x Post PII_2_

The analysis of the number of corrections on the second moment of PII (PII_2_) showed that SSG performed more corrections than SG (*p* = 0.001). The analysis of tPV (%) showed main interaction effects, *F*(2, 76) = 3.87, *p =* 0.02, *η_p_*^2^= 0.09, power (1-β) = 0.68. The post hoc detected that the tPV (%) decreased when the perturbation was inserted (PII_2_) for the SSG. Moreover, the SSG spent less tPV (%) than SG on PII_2_ (*p <* 0.05). There was difference between blocks, *F*(2, 76) = 6.15, *p =* 0.003, *η_p_*^2^= 0.13, power (1-β) = 0.87. The post hoc detected that the tPV (%) decreased when the perturbation was inserted (PII_2_) compared to Pre PII_2_ (*p <* 0.05). With th withdrawal of perturbation (Post PII_2_), the tPV (%) increased compared to PII_2_ (*p <* 0.05). There was no significant difference between groups, *F*(1, 38) = 0.47, *p =* 0.49, *η_p_*^2^= 0.01, power (1-β) = 0.10. The analysis of CE showed main interaction effects, *F*(2, 76) = 16.89, *p =* 0.01, *η_p_*^2^= 0.30, power (1-β) = 0.99. The post hoc detected that the CE increased when the perturbation was inserted (PII_2_) compared to Pre PII_2_ for SG (*p* < 0.05). With the withdrawal of perturbation (Post PII_2_), the CE decreased compared to PII_2_ for both groups (*p <* 0.05). The SSG showed similar CE in Pre PII_2_ and PII_2_ and lower CE than SG in PII_2_ (*p <* 0.05). There was difference between blocks, *F*(2, 76) = 48.37, *P =* 0.001, *η_p_*^2^= 0.56, power (1-β) = 1.00. The post hoc detected that the CE increased when the perturbation was inserted (PII_2_) compared to Pre PII_2_ (*p <* 0.05). There was no significant difference between groups, *F*(1, 38) = 3.94, *p =* 0.05, *η_p_*^2^= 0.09, power (1-β) = 0.49.

#### Pre PII_3_ x PII_3_ x Post PII_3_

The analysis of the number of corrections on the third moment of PII (PII_3_) showed that SSG performed more corrections than SG (*p* = 0.001). The analysis of tPV (%) showed interaction effects, *F*(2, 76) = 7.16, *p =* 0.001, *η_p_*^2^= 0.15, power (1-β) = 0.92. The post hoc detected that the tPV (%) decreased when the perturbation was inserted (PII_3_) for the SSG. Moreover, the SSG spent less tPV (%) than SG on PII_3_ (*p <* 0.05). There was no significant difference between groups, *F*(1, 38) = 1.23, *p =* 0.27, *η_p_*^2^= 0.03, power (1-β) = 0.19 nor blocks *F*(2, 76) = 0.94, *p =* 0.39, *η_p_*^2^= 0.02, power (1-β) = 0.20. The analysis of CE showed showed main interaction effects, *F*(2, 76) = 13.29, *p =* 0.001, *η_p_*^2^= 0.25, power (1-β) = 0.99. The post hoc detected that the CE increased when the perturbation was inserted (PII_3_) compared to Pre PII_3_ for both groups (*p* < 0.05). With the withdrawal of perturbation (Post PII_3_), the CE decreased compared to PII_3_ for both groups (*p <* 0.05). Moreover, the SSG showed lower CE than SG on PII_3_ (*p <* 0.05). There was difference between blocks, *F*(2, 76) = 95.22, *P =* 0.001, *η_p_*^2^= 0.71, power (1-β) = 1.00. The post hoc detected that the CE increased when the perturbation was inserted (PII_3_) compared to Pre PII_3_ (*p <* 0.05) and with the withdrawal of perturbation (Post PII_3_), the CE decreased compared to PII_3_. Also, there was difference between groups *F*(1, 38) = 8.82, *p =* 0.005, *η_p_*^2^= 0.18, power (1-β) = 0.82. The post hoc detected that the SSG showed lower CE than SG (*p <* 0.05).

## Discussion

In this study, we explored the impact of two discrete levels of performance stabilization: stabilization (SG) and superstabilization (SSG), on adjustments to unpredictable perturbations in intercepting a moving target. Participants were initially trained to reach one of these stabilization levels and then assigned to two groups based on their achievement. They were subsequently subjected to practice conditions featuring unpredictably timed perturbations, wherein the target’s speed was altered randomly post-movement onset across 18 random trials. The results demonstrated that superstabilization notably enhanced online adjustments in response to unpredictable perturbations and led to superior performance in challenging conditions, thus confirming our hypothesis about the benefits of superstabilization in enhancing control mechanisms and performance outcomes.

Previous implementations of the superstabilization criterion in tasks demanding isometric force control [8] and anticipatory timing tasks prior to exposure to both predictable [9, 39] and unpredictable perturbations [3] have shown enhanced outcomes under both scenarios. This is especially significant in contexts involving unpredictable perturbations, which pose substantial challenges for reorganizing action plans [4, 5]. The necessity for specialized adjustment mechanisms, such as feedback control, may clarify the benefits of superstabilization, a relationship that has not been previously explored.

Our findings indicate that superstabilization led to more frequent adjustments in scenarios involving both types of perturbations (i.e., PI and PII), as evidenced by the acceleration analysis (Fig. 2a and 3a). These adjustments entailed strategies to anticipate peak velocity closer to the movement onset, successfully implemented by the SSG in specific trials (Fig. 2b and 3b). A detailed analysis of the speed and acceleration curves showed that the strategy employed by the SSG to reduce the time to peak velocity (tPV%) enabled numerous motor adjustments and resulted in significantly improved performance accuracy (i.e., lower constant error, CE) compared to the SG in PII_2_ e PII_3_ (Fig. 2c and 3c). The necessity for a reduced peak velocity time in unpredictable environments permits individuals to employ visual feedback to modulate deceleration and implement corrections [6, 36, 40]. Such corrections were possible only for participants in the SSG, demonstrating that achieving superstabilization enhances performance variability and adaptability, thus facilitating adjustments in internal representation [9]. Moreover, superstabilization fosters more flexible behavior [25], likely due to an internal representation that bolsters the capacity to make corrections under unpredictable conditions.

Effective correction in unpredictable environments relies on adequate viewing time of the target, optimally at least 200 ms before interception [41]. If the display time is less than 200 ms, the information extracted cannot be effectively used to adjust the motor command [42], negatively impacting performance. In our experiments, participants practiced the task with movement times ranging from 200 to 250 ms. This temporal arrangement confirmed that only the achievement of superstabilization allows for the effective utilization of available time to extract relevant information, make necessary corrections, and maintain performance levels amidst perturbations.

The variances in motor control between superstabilization and stabilization are further depicted by single-subject slope curves for each level of performance stabilization. Figures 5a and 5c illustrate the velocity curves for stabilization and superstabilization, respectively, prior to Performance Increase (Pre PI), while Figures 5b and 5d display the adjustments during PI (increase in target velocity).

**Fig 5.**
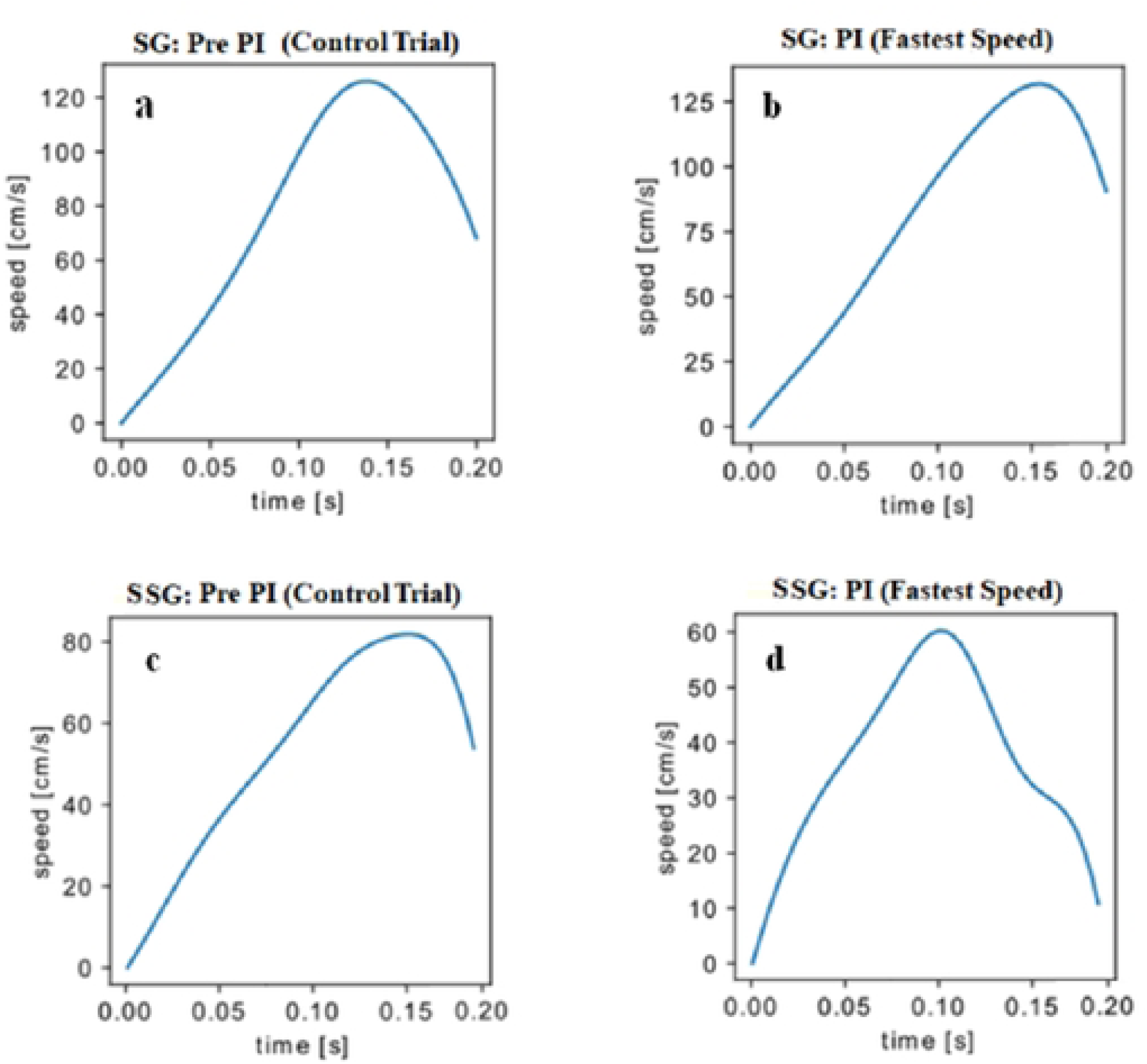
Velocity of movement (cm.s^−1^). **a.** Represents the behavior of the SG during a control trial. **b.** Represents the behavior of the SG during a trial with PI (fast velocity). **c.** Represents the behavior of the SSG during a control trial. **d.** Represents the behavior of the SSG during a trial with PI (fast velocity).

Figures 6a, 6b, 6c, and 6d display similar velocity profiles during Phase II (PII), when the target’s velocity decreases. In both Phase I (PI) and PII, the SG exhibited higher velocities compared to the SSG. Unlike the SG, the SSG anticipated the peak velocity in response to the perturbation. This anticipation gave the SSG additional time to make corrections, resulting in more accurate performance.

**Fig 6.**
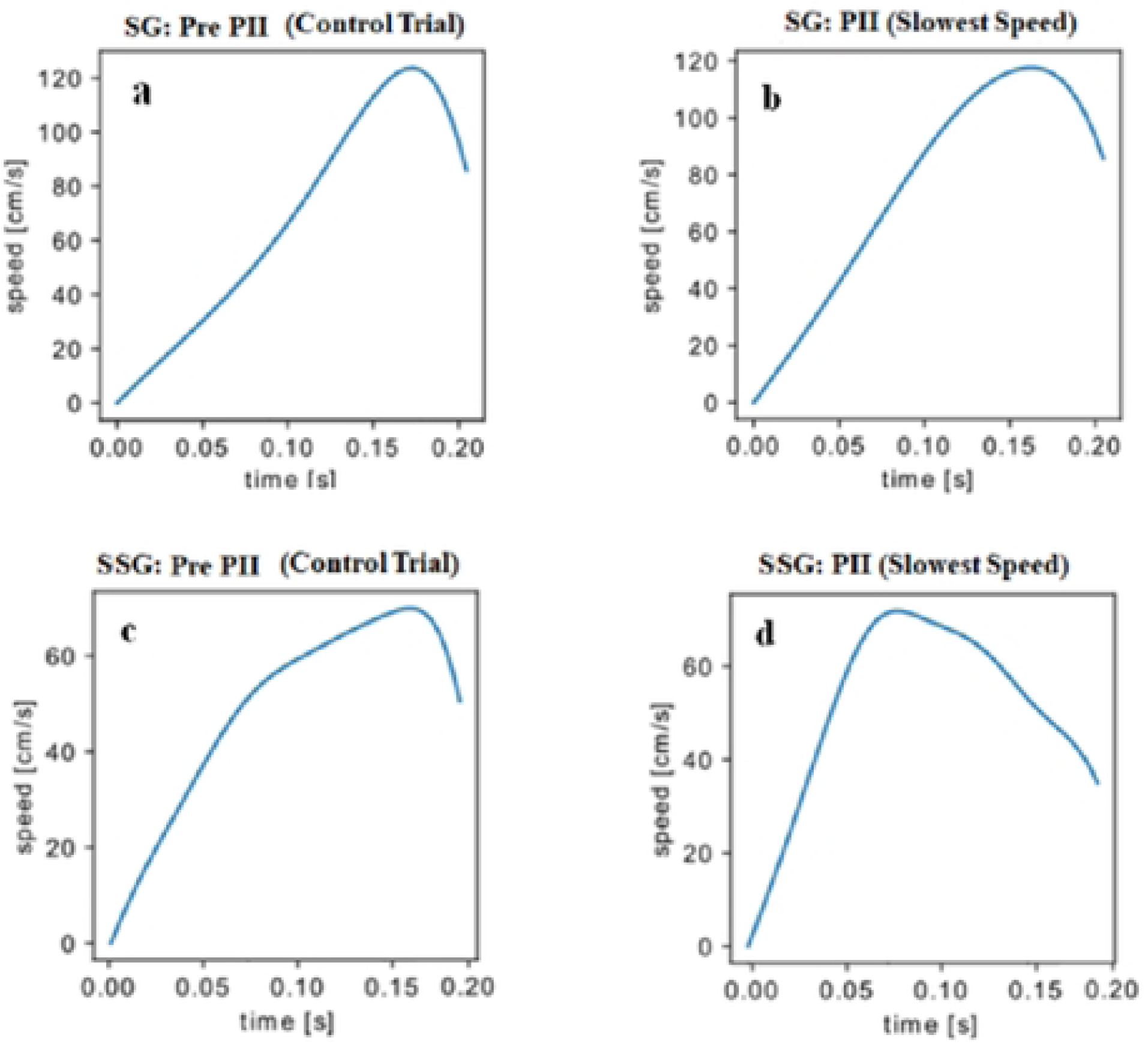
Velocity curves (cm.s^−1^). **a.** Represents the behavior of the SG during a control trial. **b.** Represents the behavior of the SG during a trial with PII (slow velocity). **c.** Represents the behavior of the SSG during a control trial. **d.** Represents the behavior of the SSG during a trial with PII (slow velocity).

Addressing unpredictable perturbations effectively requires rapid movement corrections within brief time intervals. One strategy to meet this challenge involves employing a hybrid model of motor control. In this model, a basic motor command is pre-programmed before movement initiation. Following this, the motor command is continually monitored and adjusted based on visual input, refined by a forward model embedded within the internal representation for optimal kinematic and dynamic control [43, 44]. This experiment suggests that superstabilization notably enhances the ability to extract visual information critical for responding to unpredictable perturbations. The visual system collaborates with internal models to formulate predictions [45]. Specifically, the peripheral retina and the fovea detect cues and generate error signals for rapid corrections during ongoing movement [46], as observed in our target experiment. These cues and error signals bolster the forward model, enhancing its capacity to manage the internal feedback circuit and potentially diminishing delays associated with sensory feedback in subsequent attempts [43, 45]. Such mechanisms likely account for the SSG’s superior handling of unpredictable perturbations, warranting further investigation within shorter time frames than those manipulated in the current study.

Our analysis revealed that both groups executed online corrections. However, the SSG made more adjustments than the SG, as indicated by Figures 2c and 3c data. While the SG did respond to perturbations, this group seems to have relied predominantly on information from previous attempts to control the task (i.e., pre-perturbation trials) to control the task, suggesting a less effective adjustment strategy. This reliance on previous attempts is not uncommon in interceptive tasks, such as simulated baseball batting [26], but it may lead to errors in unpredictable contexts, necessitating rapid post-detection corrections. Consequently, the SG demonstrated inefficiencies in the number of corrections needed to adjust actions, highlighting the SSG’s superiority.

In conclusion, our findings suggest that achieving a higher level of stabilization provides the sensorimotor system with more effective strategies for managing unpredictable perturbations, as evidenced by both performance and kinematic data. Moreover, these results indicate that extending practice beyond stabilization fosters a flexible internal representation of the task, capable of leveraging real-time environmental information rather than solely depending on past experiences. This insight underscores how elevated skill levels can assist performers in navigating complex motor challenges. Future investigations should further explore the impact of stabilization levels, as this knowledge has significant theoretical and practical implications across various motor learning domains, such as sports and rehabilitation, where adaptable motor behaviors are crucial.

## Limitations and Future Perspectives

We recognize that our study has limitations. The method by which we introduced perturbations enabled us to explore the effects of different performance stabilization levels in response to unpredictable perturbations, albeit limited to two magnitudes. However, predictable perturbations of varying magnitudes are commonplace in everyday life and in sports contexts. Consequently, incorporating perturbations that account for predictability and diverse magnitudes could extend our findings, providing a more comprehensive understanding of how manipulating two performance stabilization levels influences both performance and motor control.

Furthermore, we suggest exploring variations in practice schedules, such as varied practice. Research indicates that practice schedules, beyond merely the quantity of practice, can significantly affect the flexibility of internal representations [47]. This flexibility is essential for adapting effectively to perturbations, suggesting that alterations in training regimens could further elucidate the dynamics of motor control and performance adaptation. This research is being conducted in our laboratory.

## Conclusion

In line with our hypothesis, our findings demonstrate that achieving superstabilization, not merely stabilization, enhances adjustments to unpredictable perturbations and facilitates adaptation. Superstabilization appears to cultivate a more robust and flexible internal representation of the task, enabling the sensorimotor system to extract and utilize online information more effectively and make rapid movement adjustments within short time intervals.

Despite our progress in understanding motor control in tasks involving moving target interception under conditions of unpredictability and limited time for adjustments, we recommend that future studies explore manipulating the interval following the introduction of a perturbation. Reducing the time between the onset of the perturbation and its interception could challenge the ability to make adjustments using external feedback, thus necessitating reliance solely on the internal feedback mechanisms of the forward model. Such manipulation could shed light on the forward model’s capabilities when practice extends beyond basic performance stabilization.

